# Early tetrapod diversification under neutral theory

**DOI:** 10.1101/2022.02.08.476633

**Authors:** Emma M. Dunne, Samuel E. D. Thompson, Richard J. Butler, James Rosindell, Roger A. Close

## Abstract

Estimates of deep-time biodiversity typically rely on statistical methods to mitigate the impacts of sampling biases in the fossil record. However, these methods are limited by the spatial and temporal scale of the underlying data. Here we use a spatially explicit mechanistic model, based on neutral theory, to test hypotheses of early tetrapod diversity change during the late Carboniferous and early Permian, critical intervals for the diversification of vertebrate life on land. Our neutral simulations suggest, in contrast to previous studies, that increases in early tetrapod diversity were not driven by local endemism following the ‘Carboniferous Rainforest Collapse’. We show that apparent changes in face-value diversity can instead be explained by variation in sampling intensity through time. Our results further demonstrate the importance of accounting for sampling biases in analyses of the fossil record and demonstrate the vast potential of mechanistic models, including neutral models, for testing hypotheses in palaeobiology.

## Introduction

The fossil record is fundamental to our understanding of past biodiversity that can, in turn, provide important insights into future biodiversity change. The fossil record has, however, been incompletely and unevenly sampled. This has led to temporal and spatial sampling biases on estimates of deep-time biodiversity that have troubled researchers for almost half a century (Raup 1972; Smith & McGowan, 2011; Carrasco 2013; Vilhena & Smith 2013; Close *et al*. 2017; Close *et al*. 2020). A variety of different quantitative techniques, collectively known as methods of sampling standardisation, have been developed to mitigate the effects of sampling intensity biases on estimates of deep-time biodiversity (e.g., Alroy 2010; Close *et al*. 2018, and references therein). The use of sampling standardisation methods has led to significant reappraisals of the biodiversity of many fossil groups, including the first vertebrates to emerge onto land (Benson and Upchurch, 2013; Benton *et al*. 2013; Pearson *et al*. 2013; Brocklehurst *et al*. 2017, Dunne *et al*. 2018, Pardo *et al*. 2019, Brocklehurst 2020; Brocklehurst *et al*. 2020).

The establishment of terrestrial ecosystems and diversification of early tetrapods during the late Carboniferous and early Permian (323–272 million years ago) was punctuated by a climate-change-driven floral turnover at the end of the Carboniferous, referred to as the ‘Carboniferous Rainforest Collapse’ (CRC; Uhl & Cleal, 2010; Cleal *et al*. 2012). In the last decade, several studies have attempted to estimate the impact of this event on early tetrapod diversity, animals that include the earliest amphibians and reptiles. The first investigation took the early tetrapod fossil record at face-value (Sahney *et al*. 2010), largely ignoring temporal and spatial variability in sampling, and hypothesised that habitat fragmentation caused by the CRC drove increased endemism via the “island-biogeography effect” of MacArthur & Wilson (1967) which, in turn, led to a rise in global species richness coupled with a decline in local richness.

More recent investigations have questioned these conclusions on account of pervasive sampling biases in the early tetrapod fossil record and found no evidence of increases in endemism during or after the event (Brocklehurst *et al*. 2018; Dunne *et al*. 2018). Instead, Dunne *et al* (2018) found evidence of increased cosmopolitanism and lower “global” diversity following the CRC—the opposite of the pattern recovered by Sahney *et al*. (2010). The results of Dunne *et al* (2018) also indicate that fragmentation of the rainforest actually promoted the diversification of amniotes, a clade that today comprises reptiles, birds, and mammals. Still, it is apparent that sampling biases in the early tetrapod fossil record continue to mask genuine patterns of diversity and biogeography during a critical time in vertebrate evolution on land, and it is unclear to what extent these biases distort the true effects of the CRC.

Methods of sampling standardisation that attempt to mitigate the effects of sampling biases in estimates of past biodiversity have typically relied on statistical or phylogenetic techniques. Mechanistic models based on neutral theory provide an alternative approach that has not yet been widely used in palaeobiological studies (but see Holland, 2018). Mechanistic models can simulate biological communities on realistic landscapes and enable features such as palaeogeography and levels of habitat fragmentation to be experimentally manipulated. Diversity can then be sampled from the models in the same locations and to the same extent as the empirical data, thus providing a new way to test how real-world patterns of fossil record sampling impact inferred patterns of face-value (i.e., directly observed, ‘raw’, or uncorrected) diversity. As mechanistic models can be scaled up beyond the empirical sample sizes, such models can also predict wider diversity patterns, such as total species richness (gamma diversity), as well as providing estimations of detectability levels within the currently available fossil data (Brocklehurst 2015). Studies involving mechanistic models can even be used to test theories of diversity generation at global scales, much larger than could be directly perceived in the fossil record (Holland & Sclafani 2015; Jordan *et al*. 2016; Holland 2018).

Here, we test hypotheses about the impact of the CRC on early tetrapod diversity change by applying a spatially explicit variant of neutral theory to four broad scenarios of early tetrapod diversity (Figure 1). Then, we use this spatially explicit neutral model to examine the extent to which the known fossil record can infer global patterns of diversity change, thus determining how sampling biases can distort analyses of global diversity and biogeography. This study is, to the best of our knowledge, the first to apply a fully spatially explicit neutral model to empirical fossil data.

**Figure 1.**
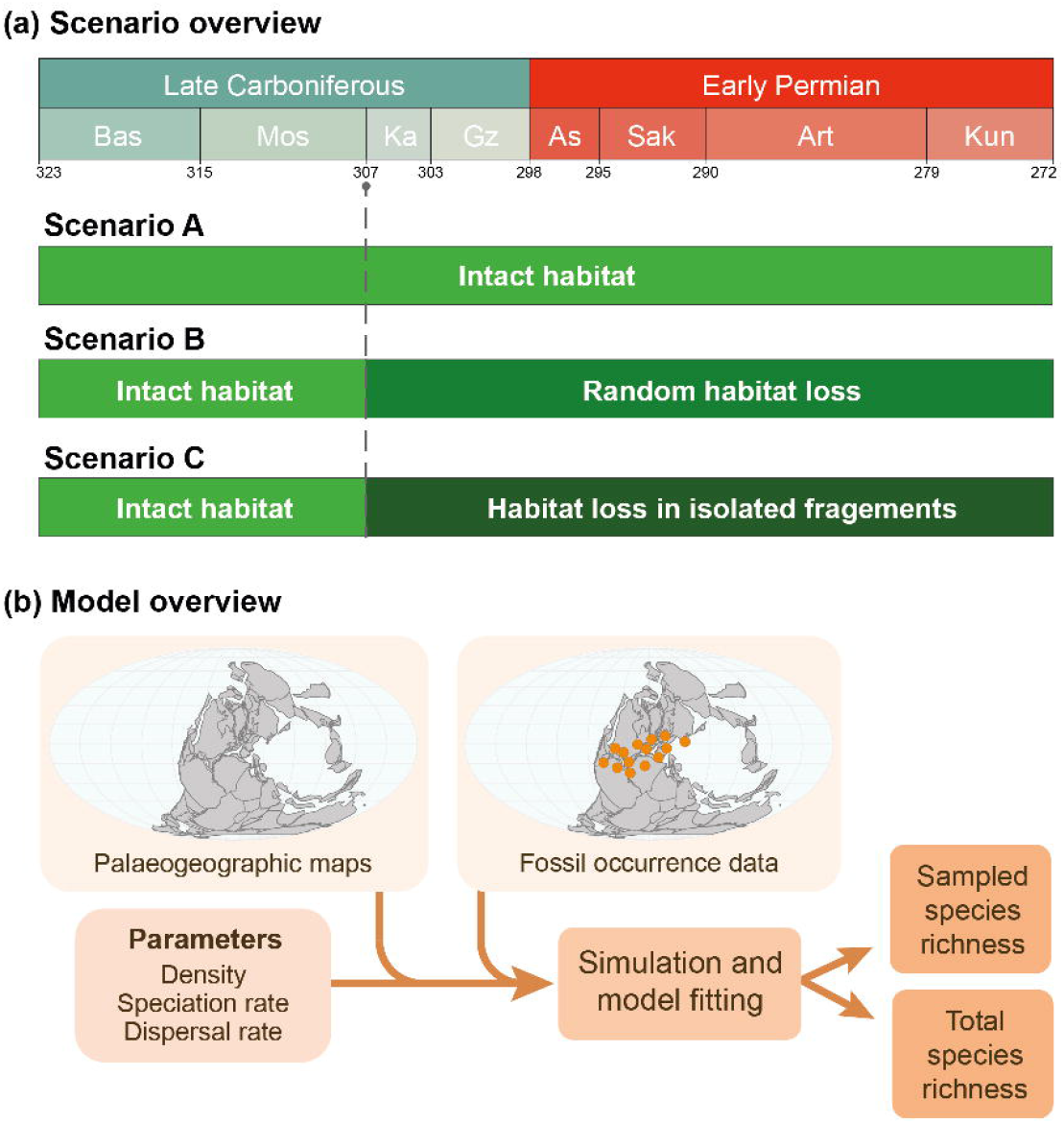
Schematic of neutral modeling of tetrapod diversity. (a) Demonstration of the simulated scenarios. Scenario A does not alter the neutral models at all due to the CRC. Scenario B models the effect of the CRC as random habitat loss across the landscape. Scenario C models the effect of the CRC as a loss in habitat such that only habitat islands remain around the localities where fossils have been found. (b) An overview of the model input data and outputs.

## Results

### Neutral models incorporating temporal and spatial sampling biases

First, simulations were performed on a pristine global landscape with no habitat fragmentation (Scenario A, Figure 1) i.e. simulations were sampled at the same palaeo-locations and to approximately the same intensity as the real fossil record. These simulated diversity patterns, which were sampled at the same palaeo-locations and to approximately the same intensity as the real fossil record, match empirical (face-value) early tetrapod diversity, as would be expected when imposing realworld sampling structure on simulations (Figure 2). From the best-fitting models and without allowing parameters to vary over time, the simulations produced 80–85% mean accuracy with the empirical fossil record calculated across four metrics of diversity (see Methods). As should be expected, the model cannot perfectly reproduce alpha, beta and gamma diversity simultaneously. When compared to the real fossil record, the neutral models of early Permian amniotes with optimized parameters produce more species (higher global richness), with slightly higher alpha diversity (Figure 1). Despite these differences, the majority of the temporal trend is reasonably well-captured for all biodiversity metrics with empirical values lying mostly within the range of variation between simulations.

**Figure 2:**
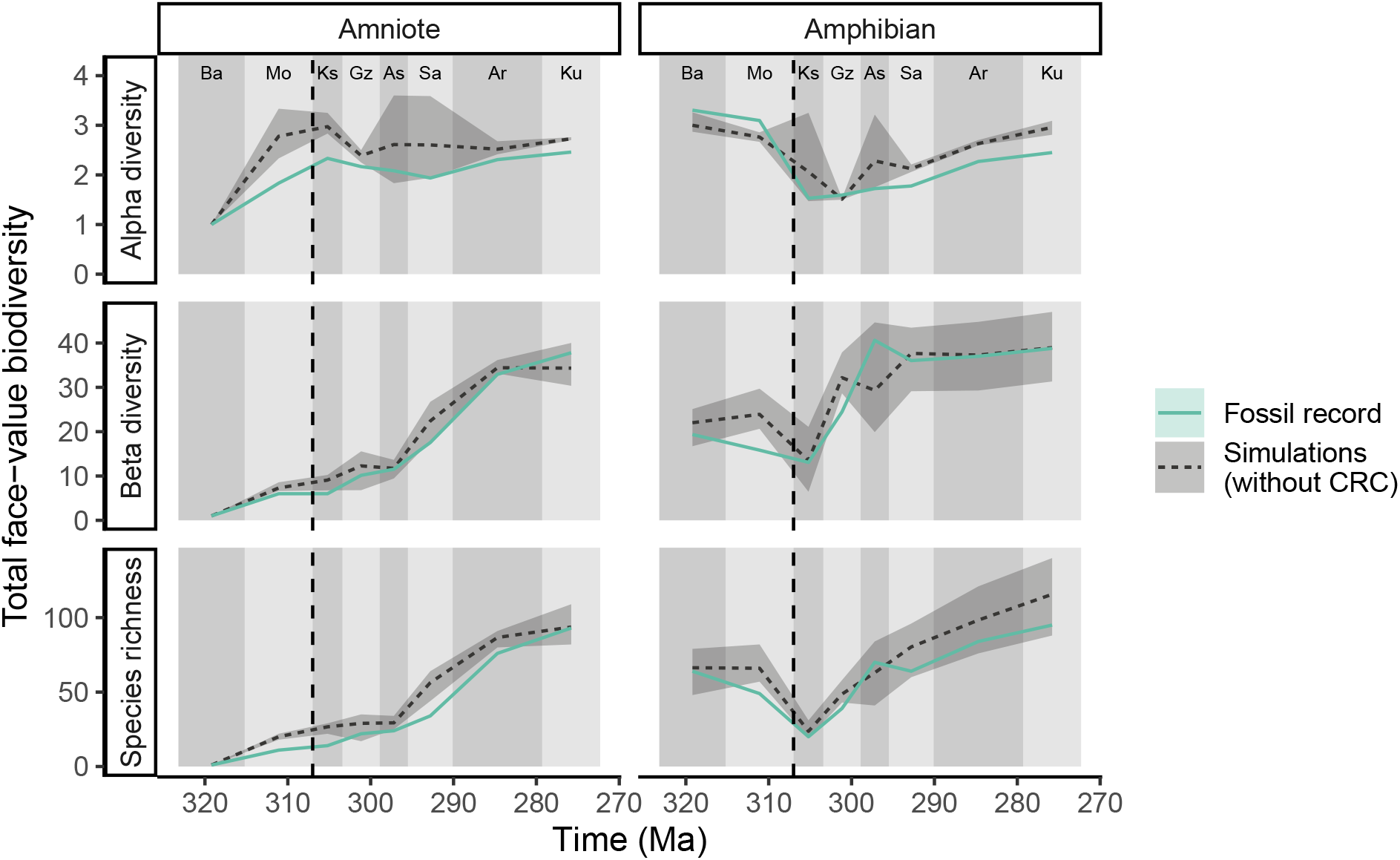
‘Pristine landscape’ (Scenario A): Simulated tetrapod diversity patterns over time compared against the fossil record (i.e. face-value, unstandardized counts of species). Alpha diversity is the mean number of species across all localities at each time point. Species richness (or “global” diversity) is the total number of species across all localities. Beta diversity is 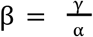. These simulations were on a pristine uniform landscape without any habitat loss or fragmentation at any point in time. Three metrics of biodiversity are shown for both amphibians and amniotes from the Bashkirian to Kungurian from empirical data (solid green line) and from simulated communities (dashed line). The shaded area represents the variation in the 5 best fitting simulations out of 25 total. The dashed vertical line indicates timing of CRC. The following abbreviations are used for time intervals shown along the horizontal axes with alternate light and dark grey shading: “Ba” = Bashkirian, “Mo” = Moscovian, “Ks” = Kasimovian, “Gz” = Gzhelian, “As” = Asselian, “Sa” = Sakmarian, “Ar” = Artinskian and “Ku” = Kungurian.

### Neutral models under habitat fragmentation

To test the hypothesis that fragmentation of the rainforest at the end of the Carboniferous promoted the development of endemism amongst early tetrapod communities, we modelled two scenarios of habitat loss and fragmentation from 307 Ma onwards: habitat loss in a random pattern (Scenario B, Figure 1) and habitat loss in a clustered pattern (Scenario C, Figure 1). The random habitat scenario maintains connectivity across the landscape but can still contain considerable habitat loss. The clustered habitat scenario leaves isolated islands of habitat that will promote endemism within each island over geological timescales. This simulates a scenario where the islands of habitat correspond conceptually to islands from MacArthur and Wilson’s theory of island biogeography (1967), thus directly testing the theory of Sahney *et al*. (2010) that endemism, driven by fragmentation, is the cause of tetrapod diversity increases post-CRC.

Our models of random habitat loss (Scenario B) demonstrate that increasing the amount of habitat loss, while keeping all other parameters the same, causes “global” species richness and beta diversity to decline (Figure 3, Supplementary Figure 5). Species richness decreases relatively linearly across all time periods. However, alpha and beta diversity demonstrate a more variable pattern across time for different levels of habitat loss. In particular, the interval between 307–297 Ma has very similar alpha diversity for all levels of habitat loss, potentially caused by the lower numbers of fossils found at this time (Supplementary Figure 2), as alpha diversity will then be impacted primarily by the number of individuals, rather than the surrounding habitat.

**Figure 3:**
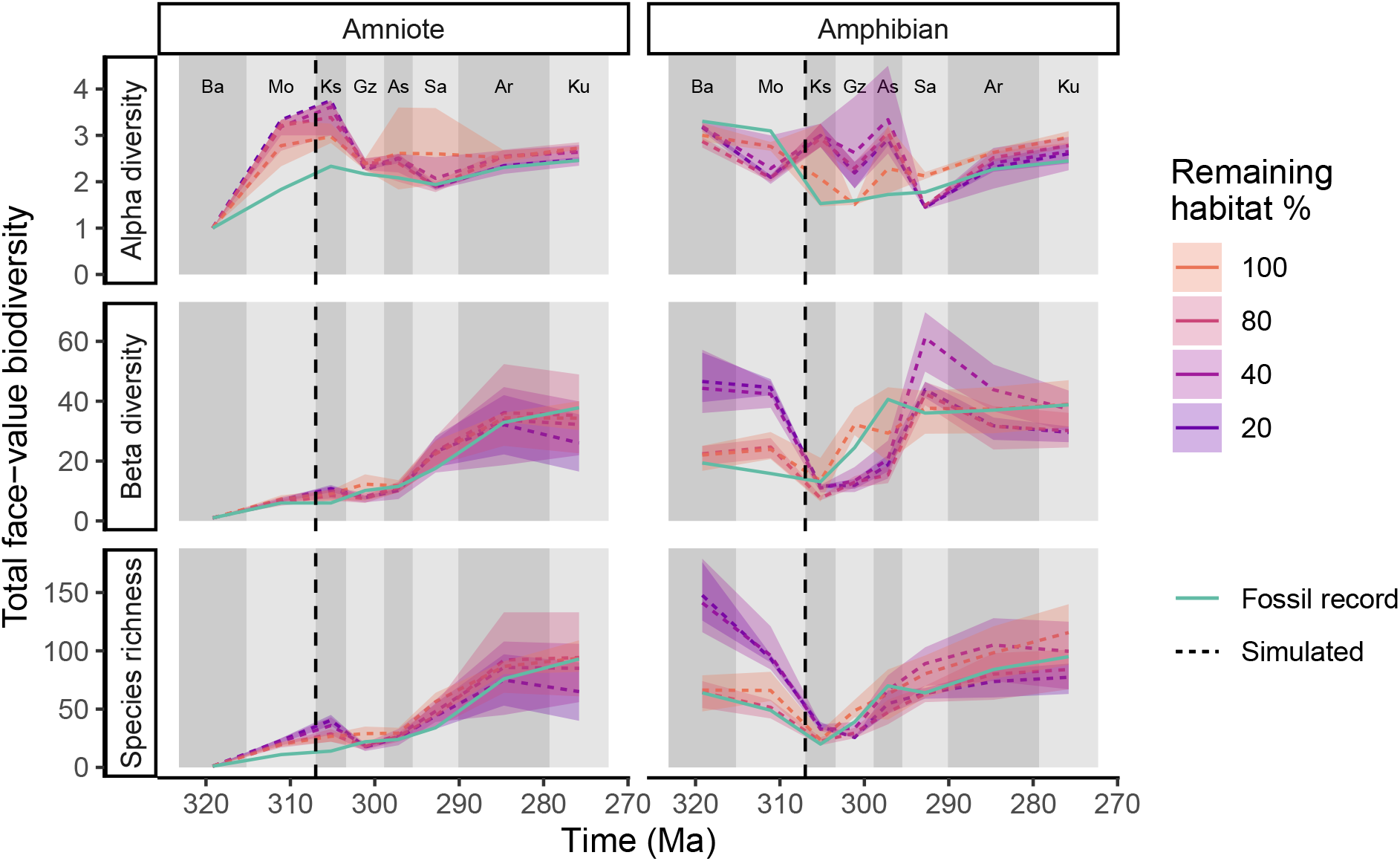
Random habitat loss (Scenario B): Simulated tetrapod diversity patterns over time compared against the fossil record (i.e. face-value, unstandardized counts of species). Alpha diversity is the mean number of species across all localities at each time point. Species richness is the total number of species across all localities. Beta diversity is 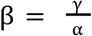. Here, the impact of the CRC is represented by random habitat loss occurring at 307 Ma (dashed line). The same set of parameters is shown for four degrees of habitat loss (from no habitat loss, i.e. 100% remaining, to 80% loss, i.e. 20% remaining). Additional results for 20, 40 or 80% habitat remaining are given in Supplementary Figure 5. Interval abbreviations are as in Figure 2.

Our clustered habitat scenario C tested whether neutral theory supports the hypothesis that habitat loss results in highly disconnected habitat islands that promote endemism. Under these circumstances, unless the fossil localities were close, dispersal between distinct fossil localities was almost entirely restricted, meaning that the number of shared species between localities was likely to be very low. The neutral simulations of the clustered habitat scenario generated diversity patterns that did not closely fit the empirical fossil data (Figure 4). Although the overall trend matches to some extent, the simulations had a high level of variability between intervals, primarily dictated by the number of fossil localities known for each interval. Furthermore, the loss of habitat and resulting decrease in size of the metacommunity supplying individuals to the fossil sites caused a reduction in species richness. Similarly, there was also a reduction in alpha diversity, particularly for amniotes.

**Figure 4:**
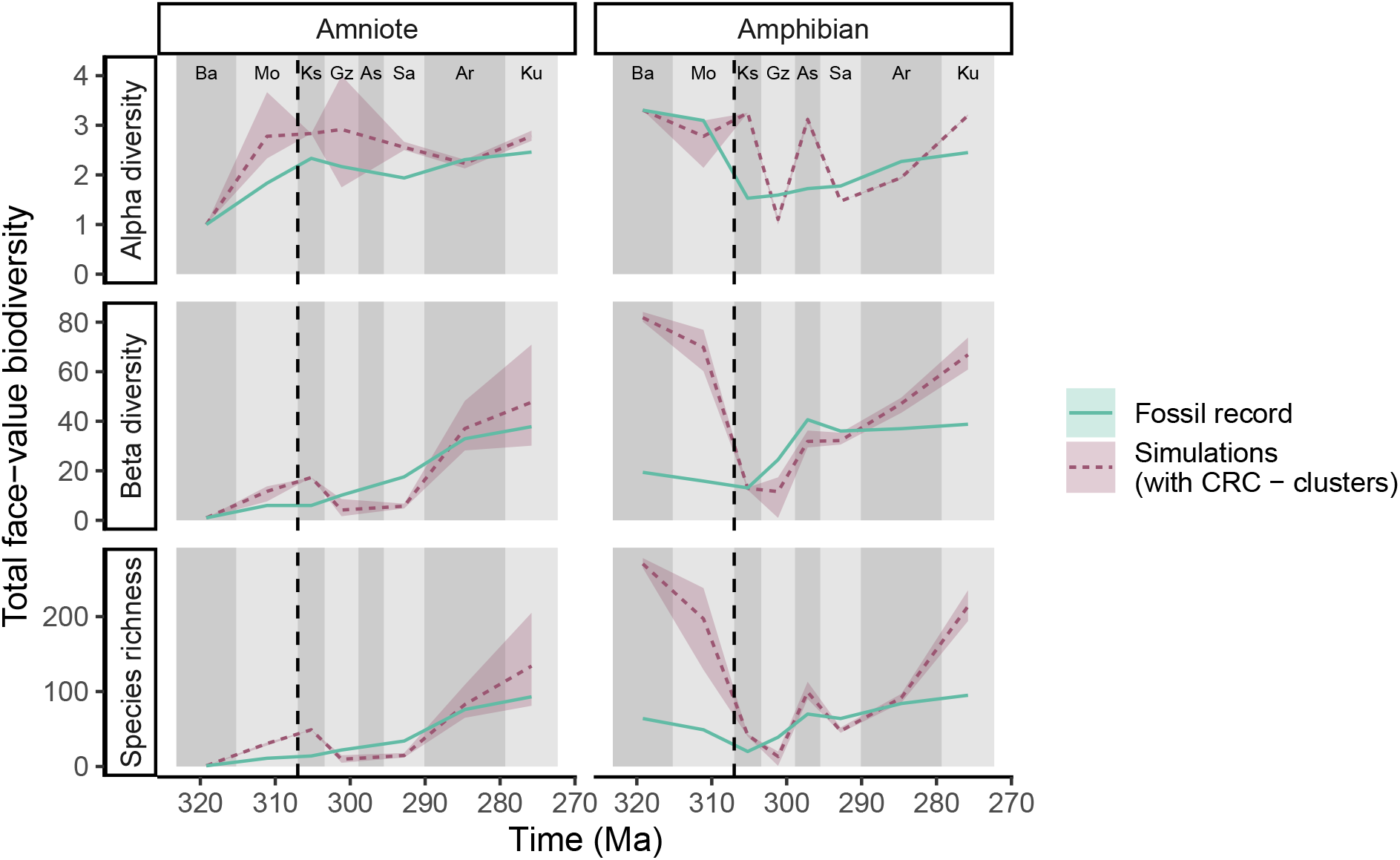
Clustered habitat scenario (Scenario C): Simulated tetrapod diversity over time from neutral models compared against the fossil record (i.e. face-value, unstandardized counts of species). Alpha diversity is the mean number of species across all localities at each time point. Species richness is the total number of species across all localities. Beta diversity is 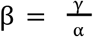. Neutral simulations incorporated habitat loss from the CRC following the clustered habitat scenario. The remaining habitat following the CRC is associated into habitat islands 100km diameter at the locations of each fossil locality. The shaded area represents the variation in the five best fitting simulations. The dashed line at 307 Ma indicates the timing of the CRC. Interval abbreviations are as in Figure 2.

By simulating this same best-fitting scenario (20% random habitat loss for amniotes and a pristine landscape for amphibians), but sampling more individuals at each locality, it is possible to simulate the broader diversity changes under the same model. When ten times more individuals are sampled from each fossil locality (Figure 5), differences emerge when compared with simulations where the fossil record is exactly matched. The general trends in species richness over time for both amniotes and amphibians are roughly similar to the trends observed in the fossil record (Figure 5). However, there is no longer a significant increase in beta diversity post-CRC, especially for amphibians. Likewise, alpha diversity is relatively consistent over time. There is also a broader range in the simulation outcomes once more individuals are sampled.

**Figure 5:**
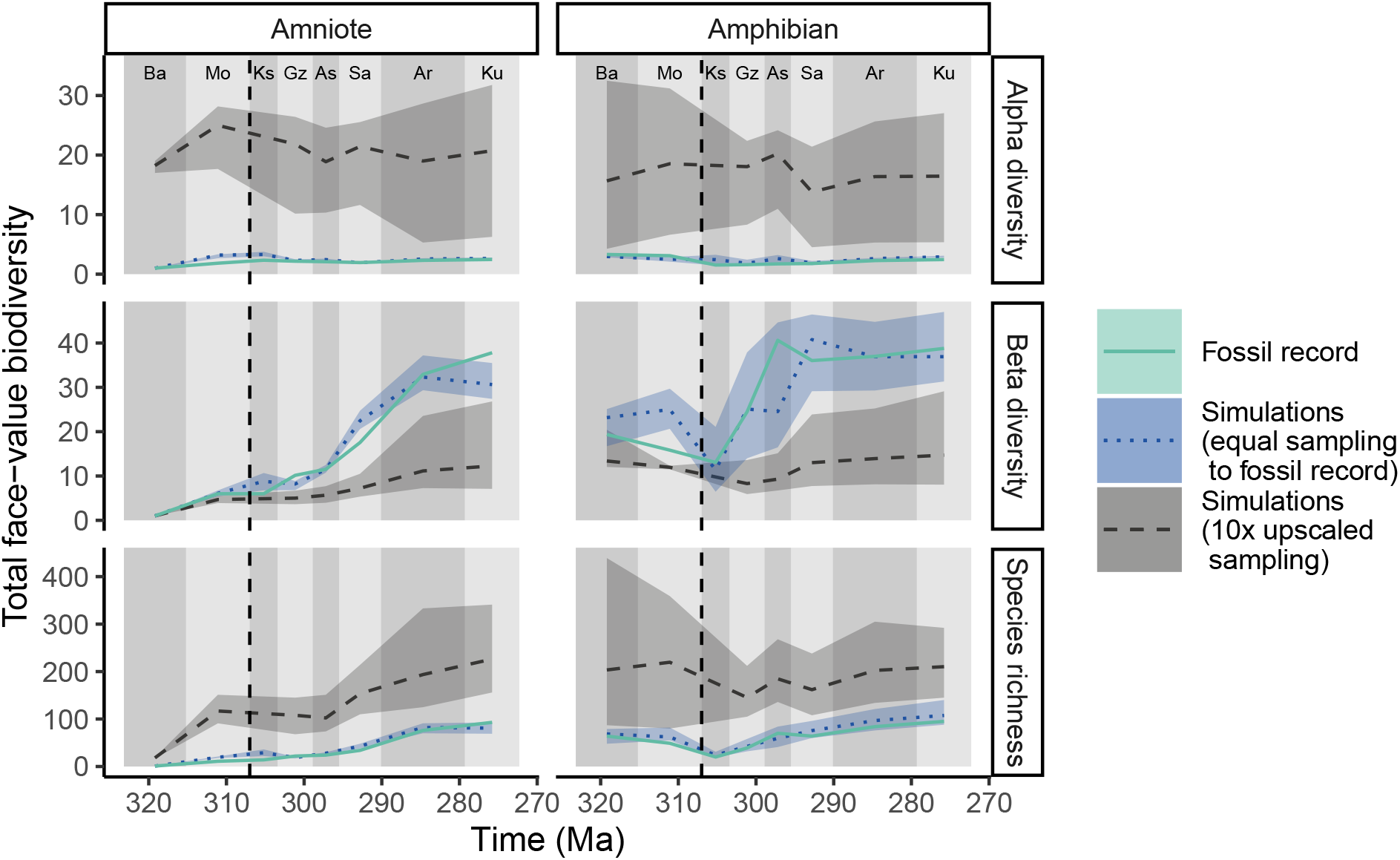
‘Upscaled diversity’ from the fossil record using neutral models. Solid green lines represent the empirical data from the fossil record. Blue dotted lines represent the mean values from simulations of the five best-fitting parameters from the scenario under random clearing (as per Figure 3). The grey dashed lines represent the same simulations, but sampling ten times more individuals than is present in the fossil record. The shaded areas represent the variation in the five best fitting simulations. The dashed line at 307 Ma indicates the timing of the CRC. Interval abbreviations are as in Figure 2.

To remove temporal variation in sampling intensity (but retain spatial sampling structure), a scenario was simulated with even sampling effort over time. When 100 individuals are randomly selected from each time slice, in the same spatial arrangement as the empirically sampled localities, the trend in species richness over time closely follows the changes in global density (Figure 6). The simulated patterns in diversity where sampling effort is standardised bears only limited resemblance to the real fossil record together with its sampling biases, matching the general trend only for beta diversity.

**Figure 6:**
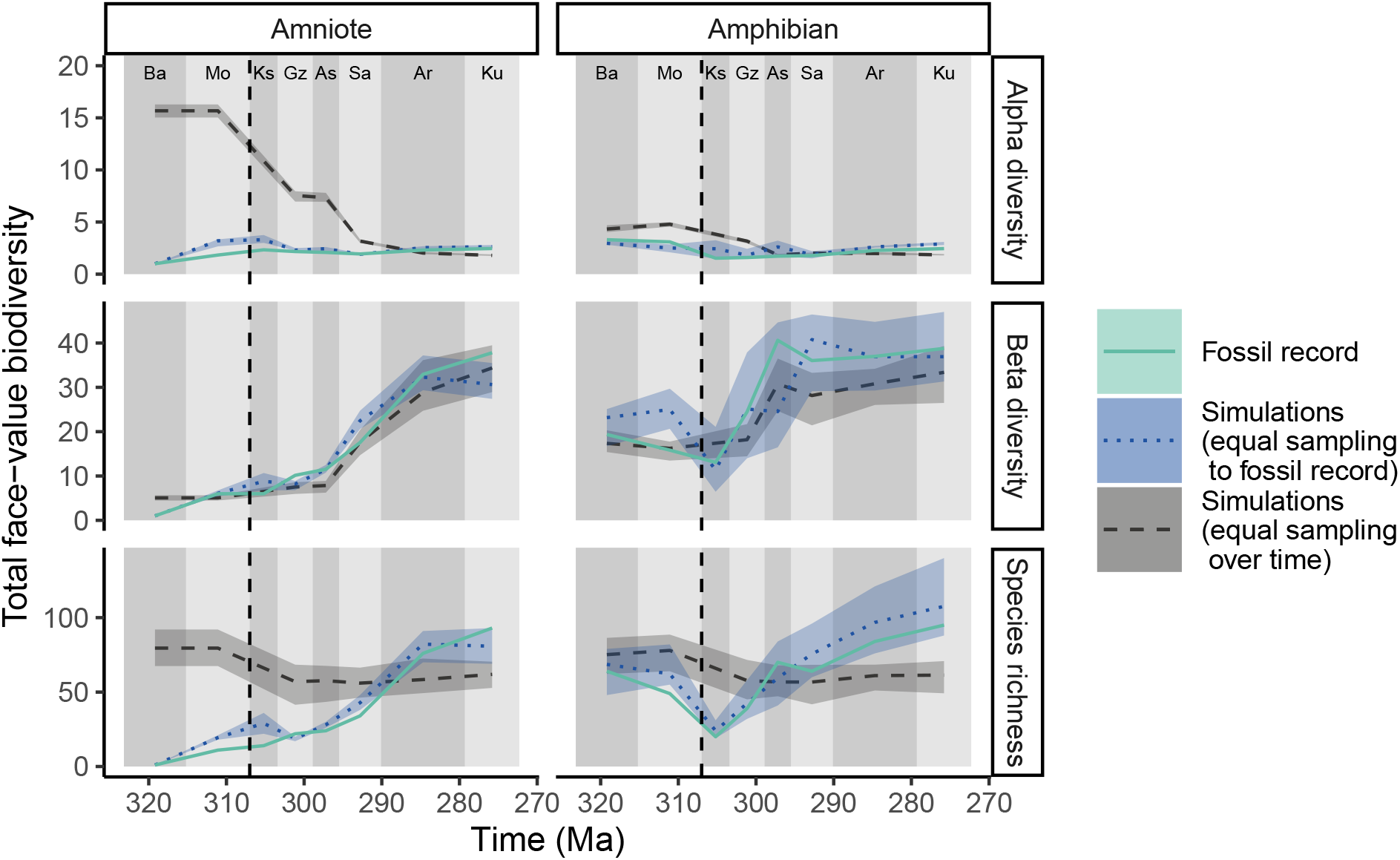
Tetrapod diversity from neutral models where temporal sampling biases are removed but spatial structure of sampling is retained. Green solid lines represent the empirical data from the fossil record. Blue dotted lines represent the mean values from simulations of the five best-fitting parameters from the scenario under random clearing (as per Figure 3). The grey dashed lines represent the same simulations, but randomly sampling 100 individuals for each interval. These simulations are without any temporal bias in sampling frequency in contrast to the fossil record and preceding simulations where the number of individuals sampled changes through time. The shaded areas represent the variation in the five best fitting simulations. The dashed line at 307 Ma indicates the timing of the CRC. Interval abbreviations are as in Figure 2.

## Discussion

Based on our neutral models, apparent increases in face-value diversity observed in the fossil record can be explained by a simple mechanistic model that accounts for biases in sampling. However, there does appear to be a small but observable change in the characteristics of early tetrapod diversity around 307 Ma, the approximate timing of the ‘Carboniferous Rainforest Collapse’. This can either be explained by changes in dispersal, changes in species density (Supplementary Figure 3-4), or fragmentation of habitats (which is theoretically similar to a reduction in species diversity, Figure 3). These findings support the previous assessment that patterns of diversity in the early tetrapod fossil record should not be interpreted at face-value (Dunne *et al*. 2018).

The model scenario of rainforest fragmentation that is most consistent with the empirical (face-value) fossil data is one where the global density of early tetrapods decreases by a small amount at 307 Ma (Figure 3). When sampling the simulations in a realistic manner, this results in a decrease in total species richness and beta diversity. Under this scenario, total diversity losses during the Carboniferous Rainforest Collapse are even greater than observed in the fossil record (after accounting for the changes in sampling effort over time). The development of endemism is not enough to offset the diversity decrease from habitat loss.

When the same models were simulated, but many more individuals sampled, the emergent diversity patterns changed considerably (Figure 5). This finding suggests that the temporal changes in alpha and beta diversity found in the fossil record may disappear as more fossils are found, reducing the impact of sampling biases. Furthermore, when the same number of individuals are sampled from each point in time within our models, the trends in species richness and alpha diversity mostly disappear (Figure 6). This, again, suggests that the face-value patterns in the fossil record are an artifact of changes in the number of locations sampled within each interval. The spatial coverage of the fossil record for early tetrapods falls a long way short of being truly global—and indeed varies through the intervals of geological time being studied.

Taken together, our results suggest that endemism from habitat loss at the CRC led to a net decrease in biodiversity and not an increase as has been previously claimed Sahney *et* al (2010),. After accounting for sampling bias, the limited changes to global richness are primarily driven by a minor reduction in global early tetrapod density over time, which is consistent with the expected ecological impact of the collapse of the rainforests and drying of the climate. The simulated scenario that aligns best with the empirical, face-value patterns is one of random habitat loss of roughly 0-20%, a scenario that is dynamically identical under neutral theory to an equivalent reduction in density (Thompson *et al*., 2019).

Our models used relatively abstract patterns of habitat loss, as the real patterns are not known. Future research could attempt to produce more realistic patterns of rainforest habitat loss, based on either palaeoclimate reconstructions or comprehensive occurrence data for fossil plants. Integrating more accurate maps of tropical rainforest coverage over time with the mechanistic basis of neutral theory would be more informative for exploring theories of diversity generation following the CRC. This is not currently possible due to the absence of readily available palaeoclimate reconstructions for this particular time interval and the lack of a comprehensive, spatially explicit, occurrence-based database for fossil plants, similar in structure and content to the Paleobiology Database. It is not immediately clear how one would relate forest patterns to the dynamics of early tetrapod diversity, as amphibians and reptiles (both modern and extinct) exhibit broad variability in their dependency on forest cover. One immediate pattern of rainforest loss that could be incorporated into future related work, with the addition of empirical data, is the hypothesis that the rainforest disappearance began in western Pangaea before moving eastwards (Cleal & Thomas 1999, 2005).

The neutral models explored here assumed that densities were consistent over time, except in the case of habitat loss at the CRC. However, the abundance (population density) of early tetrapods would also have a significant effect on the numbers of specimens preserved in the fossil record. Consequently, lower numbers of fossil specimens could be indicative of smaller populations and lower species richness. Yet it is difficult to truly resolve the relationship between density and sampling rate because the nature of fossil preservation varies substantially over time. Across our dataset of late Carboniferous and early Permian tetrapods, quality of preservation (and thus the size of the “taphonomic window”) varies substantially, which in turn influences sampling intensity. Fossil localities of late Carboniferous age that have yielded particularly well-preserved or abundant specimens are typically coal deposits (e.g. coal mines at Nyrañy in the Czech Republic and Linton Diamond Mine in Ohio, USA). In the early Permian, due to the combination of orogenic activity and drier climatic conditions, fossils are much less likely to be preserved in coal deposits. Instead, many richly diverse localities in the early Permian are located in sandstone quarries that have been extensively excavated over many decades (e.g. various localities in the Red Beds of Texas and Oklahoma, USA). Due to these temporal changes in preservation it is impossible to infer true densities of early tetrapods during this interval (and likely any interval in the geological past). This limitation motivated keeping density as a free parameter within the neutral simulations but precludes understanding of how both early tetrapod densities and preservation rates varied. The resolution of this problem requires a more intimate understanding of both the true densities of early tetrapods over time and changes to the preservation rates over time (one of the measures that is possible to estimate for species within assemblages).

Our explanations of the changes in early tetrapod diversity through time have all been based on ecological neutral theory. Alternative explanations could come from changes in non-neutral dynamics, such as species niche structure, competition between species, or wider ecosystem-level shifts. These explanations cannot yet be tested from a mechanistic basis but represent an exciting avenue of future research.

## Conclusions

Statistical approaches can provide important insights into patterns of diversity (Lloyd, 2012; Mannion *et al*., 2012; Close *et al*. 2018; Dunne *et al*. 2018), however, they are generally limited by the geographical and temporal extent of the available fossil occurrence data. In our study, spatially explicit neutral models have proven to be a valuable tool for directly testing established hypotheses of diversity change in the first vertebrates to emerge onto land, and illuminating the impacts of spatial and temporal sampling biases on their face-value diversity patterns.

The approach we used for testing hypotheses of diversity in the fossil record represents a novel direction for integrating modern ecological theory with palaeontological data. Such interdisciplinary studies have been identified as crucial for informing predictions for future diversity (Benton 2016; Barnosky *et al*. 2017) as well as more accurately understanding past biodiversity patterns (Jackson 2002; Willis & Birks 2006; Bonuso 2007; Mayhew *et al*. 2008). Our results shed further light on the impact of the CRC on early tetrapod diversity, by showing that increased endemism resulting from habitat loss at the CRC is unlikely to have produced an increase in biodiversity. Our study also offers new insights into the effects of sampling bias on fossil diversity estimates, and demonstrates the huge untapped potential that mechanistic models, such as those founded on neutral theory, have for testing hypotheses of deep-time biodiversity change.

## Methods

### Neutral models

Assessment of spatial and temporal biases on a mechanistic basis by definition requires a model that is spatially and temporally explicit. In addition, to study the impact of habitat loss and fragmentation on biodiversity requires a model that can directly incorporate these dynamics within the biodiversity-generating process. Neutral models fulfil all these requirements and are also tractable at large scales. Neutral theory (Hubbell 2001) assumes that the properties of an individual are independent of its species identity. The dynamics of neutral models are thus dictated by some combination of dispersal, ecological drift and speciation. The output of neutral models is a simulated ecological community, where each individual has an assigned species identity. These simulated communities are equivalent to a complete census of the simulated area. The communities provide a baseline for expected biodiversity under “idealised” conditions (Alonso *et al*. 2006), against which the biodiversity from real communities can be compared. Neutral theory has, however, only rarely been applied in analyses of fossil data. A few palaeoecological studies have used spatially implicit neutral theory (Holland & Sclafani 2015; Jordan *et al*. 2016; Holland 2018) where populations (e.g. within separate continents) are divided to roughly represent spatial barriers. To the best of our knowledge, however, no previous study has applied a fully spatially explicit neutral model to fossil data.

The classic, spatially implicit model (Hubbell 2001) conceives of a local community connected to a metacommunity by immigration at a given rate, other models incorporate more explicit dispersal between parts of the landscape. Being based on fundamental biological mechanisms, neutral theory has utility for identifying underlying dynamics (Vergnon *et al*. 2009), acting as a null or “ideal” model (Alonso *et al*. 2006) or making predictions at broader spatial or temporal scales than are possible with field experiments (Rahbek *et al*. 2007). We use a spatially explicit neutral model (Rosindell & Cornell 2007) that incorporates the exact locations of each individual in space and incorporates a dispersal kernel to describe the distance moved by offspring from their parents. Such a fully spatially explicit model is essential to account for spatial sampling bias. The metacommunity concept of the spatially implicit model is replaced by movement around a broad spatially explicit landscape.

The mechanism of our model proceeds as follows, an individual is first chosen to die leaving a ‘space’ that will be filled by a new-born individual. The parent of the newborn individual is chosen from other nearby individuals according to a dispersal kernel, which we modelled as a two-dimensional normal distribution. The new-born is normally conspecific to its parent, but occasionally, with probability *v* at each birth, it becomes a new species. Over many generations, nearby individuals are more likely to be the same species, whereas distant individuals will be more likely to be different. We use these models to generate communities of species across the landscape.

A major development for neutral theory was backwards-time coalescence methods (Rosindell *et al*. 2008), which produce equivalent results to a naïve (forwards-time) implementation of the mechanisms described above but are many orders of magnitude faster in computational performance. Furthermore, many scenarios are made possible with coalescence that are not possible otherwise, such as exceedingly large or infinite landscapes (Rosindell & Cornell 2007) or sampling a small subset of individuals from the landscape without having to simulate the entire landscape first. The latter feature means that our models can simulate observations at just the precise locations observed in the fossil record, whilst accounting mechanistically for the whole community alive at the time with a full spatial structure from the relevant period in history. An equivalent model using forwards-time techniques would require simulating every tetrapod that existed across the entire time frame and continent of interest, a feat not remotely feasible with current computational power. Unfortunately, almost all non-neutral models cannot benefit from the use of coalescence and associated abilities to account for sampling in huge spatially explicit systems. We use the pycoalescence package available for Python and R (Thompson *et al*, 2020), which uses coalescence methods implemented in C++ for high performance spatially explicit neutral simulations. All simulations were performed on high-throughput computing systems at Imperial College London.

### Preparation of fossil occurrence data

Data detailing the global occurrences of early tetrapod species from the late Carboniferous (Bashkirian) to early Permian (Kungurian) were downloaded from the Paleobiology Database (www.paleobiodb.org). These data represent the published knowledge on the global occurrences of early tetrapod species alongside taxonomic opinions; it is the result of a concerted effort to document the Palaeozoic terrestrial tetrapod fossil record. The dataset was cleaned by removing marine taxa, ichnotaxa, and taxa with uncertain taxonomic identifications. The total number of amniote (including Reptiliomorpha) and amphibian (i.e. non-amniotes and early tetrapodomorphs) species per locality was ascertained and recorded. The resulting dataset (see supplementary material) details the number of amniote and amphibian species found at each locality (i.e. a “collection” in PBDB terms) during each of the eight stratigraphic intervals from the Bashkirian to the Kungurian.

### Neutral simulations of early tetrapods

We split the tetrapods into amphibians and amniotes to reflect their differing physiologies and environmental preferences, treating each with an independent neutral model. Our simulations required maps of the relative density of individuals across the globe. These were determined separately for each interval (Bashkirian–Kungurian) from the continental boundaries of the time. Global rasterized maps of individual relative densities were produced at 0.01-degree resolution using the continental extents provided by the Paleobiology Database based on GPlates palaeogeographical reconstructions (Seton *et al*. 2012). This corresponds to pixels of around 1 km^2^ each representing a cell for our model. The palaeocoordinates of each fossil locality were calculated and localities were then aggregated within each 1 km^2^ cell. Specimen counts per locality were estimated using the ‘occurrences-squared’ heuristic (Alroy 2000), calculated simply as the square of the number of unique fossil occurrences. This metric provides a basic way of accounting for the fact that most localities lack information about counts of specimens, and because it is rarely obvious how many distinct individuals contributed to a set of fossil fragments. Using this metric in our models approximates the total number of individuals that contributed to the observed fossil record and therefore the number of individuals that should be sampled in the neutral simulations. This generated a ‘sample map’ defining the number of individuals to be sampled at each position in space. As the majority of the globe was not sampled, most cells in this sample map were set to 0. The relative density and the sample maps together contain the spatial information of the entire global community of amphibians and amniotes for the simulation and define which individuals from each global community were sampled.

The second parameter critical for the simulations is the dispersal rate (σ), which controls the distance that individuals disperse across the landscape in a given generation. σ is used as the variance in a Rayleigh distribution determining the radius of dispersal, with a separate uniform random number determining the direction of dispersal. This means that larger values of σ correspond with longer dispersal distances, on average.

The intervals sampled from the fossil record were sufficiently far apart in time that we reasonably assumed no shared species between the different time intervals within the model. Consequently, we ran simulations for each time interval as separate neutral models in parallel, and aggregated the communities post-simulation.

We performed simulations with parameters encompassing a broad range of biologically-feasible values: density values for habitat cells ranged from 25–1000 individuals per km for “habitat” regions (non-habitat regions have a density of 0 individuals), the parameter of dispersal (σ) varied to give mean distances of 0.1–14km, and speciation rates varied from 10^−8^ to 10^−1^. We explored 5 density and 5 dispersal parameters giving 25 combinations using Latin hypercube sampling (McKay *et al*. 1979) to evenly sample from arithmetic parameter space. Under coalescence methods, higher speciation rates can be applied post-simulation for generating communities (Rosindell *et al*. 2008; Thompson *et al*. 2020). We performed simulations using a minimum speciation rate of and applied all other speciation rates afterwards to generate additional communities.

Four broad scenarios of tetrapod diversity were simulated (Figure 1). In all models, the global landscape was restricted by continental boundaries. Our simplest model (Scenario A) contained pristine habitat with no habitat loss (i.e. uniform, with no habitat fragmentation) Two scenarios (Scenarios B and C) exhibited habitat loss of different forms following the CRC. The landscape was fragmented according to a random spatial pattern, so that land areas contained habitat on a percentage of their area (either 20%, 40% or 80% of habitat remaining). The random pattern was generated by randomly removing pixels from the landscape until the desired percentage of habitat remains.

### Model parameterisation

In order to determine how well the simulations fit the patterns in the fossil record, four biodiversity metrics were used for each interval: the alpha diversity (α) for each fossil locality (i.e. the local species richness), the mean alpha diversity across all localities, the total species richness across all localities (γ), and the mean beta diversity (calculated as 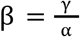) across all localities. The mean actual percentage error between the real and simulated fossil records in alpha diversity for each locality was averaged to get a mean alpha accuracy μ_α_. The mean actual percentage error between the real and simulated fossil records was calculated for each other metric (α, β and γ). Averaging the mean actual percentage errors for the four metrics (μ_α_, α, β, and γ) gives an indication of the goodness of fit for one simulation–we refer to this percentage as the accuracy of a single simulation. There is some redundancy between the values as the parameters are not independent, but the approach should still result in the simulation that most closely matches the real fossil record.

As each interval was run as a separate neutral simulation, the parameters of speciation rate, density and dispersal could be allowed to vary over time. However, as combinations of parameters can be aggregated in any number of ways, we considered just two possibilities that reflected our assumptions of the possible ecological changes over time: either there was no change in these parameters (i.e. we use a single parameter set for all intervals), or the parameters could change at the time of the CRC (i.e. we use two parameter sets - one for pre-CRC [323–307 Ma] and one for post-CRC [307–372 Ma]). The first scenario represents a neutral ecosystem with no changes in fundamental ecological dynamics. The second presents a neutral scenario that assumes ecological changes were generated by the CRC and may be reflected in neutral dynamics. In some tests a single set of parameters (speciation rate, dispersal and density) was used for all time intervals, whilst in others, this requirement was relaxed to investigate how the parameters themselves may change over time.

### Upscaling and downscaling simulated communities

In order to explore potential biodiversity patterns that would emerge if the fossil record included a larger number of individuals, we ran simulations with the same model parameters as the best-fitting simulations, but sampled ten times more individuals. This scenario demonstrates how the emergent biodiversity patterns change with the proportion of individuals captured in the fossil record. Conversely, we also explored the effect of sampling the same number of individuals from across intervals, rather than following the temporal changes in sampling intensity present in the fossil record. Doing so samples from the simulation without sampling-intensity biases but retains the spatial biases of the real world sampling pattern.

## Data and code Availability Statement

All relevant data and code supporting our analyses are available at: https://github.com/thompsonsed/palaeo_neutral_sims

## Acknowledgments

We thank all contributors to the Paleobiology Database, in particular T. Liebrecht, R. Whatley, J. Dummasch, J. Alroy and M. Carrano. This is Paleobiology Database official publication number **XXX**. EMD would like to thank T. Dunkley-Jones and P. Mannion for helpful comments and discussion. EMD, RAC, and RJB were funded by the European Union’s Horizon 2020 research and innovation programme under grant agreement 637483 (ERC Starting Grant TERRA to RJB), and also through a Leverhulme Research Project Grant (RPG-2019-365 to RJB, Sarah Greene, Roger Benson and Dan Lunt). SEDT was funded by the Imperial-NUS Joint PhD Scholarship. J.R. was funded by a NERC fellowship (NE/L011611/1). Through JR and SEDT, this study is an output of the Georgina Mace centre for the Living Planet at Imperial College London.

## Author contributions

R.A.C and J.R. conceived the project and all authors input into the design. E.M.D. and S.E.D.T curated the data, conducted the analyses, prepared figures, and led the writing of the manuscript. All authors contributed to collating material for the supplementary information, and to the writing and approval of the final manuscript.

