## Supplementary figures and images for "Early tetrapod diversification under neutral theory"

### Supplementary Figure 1

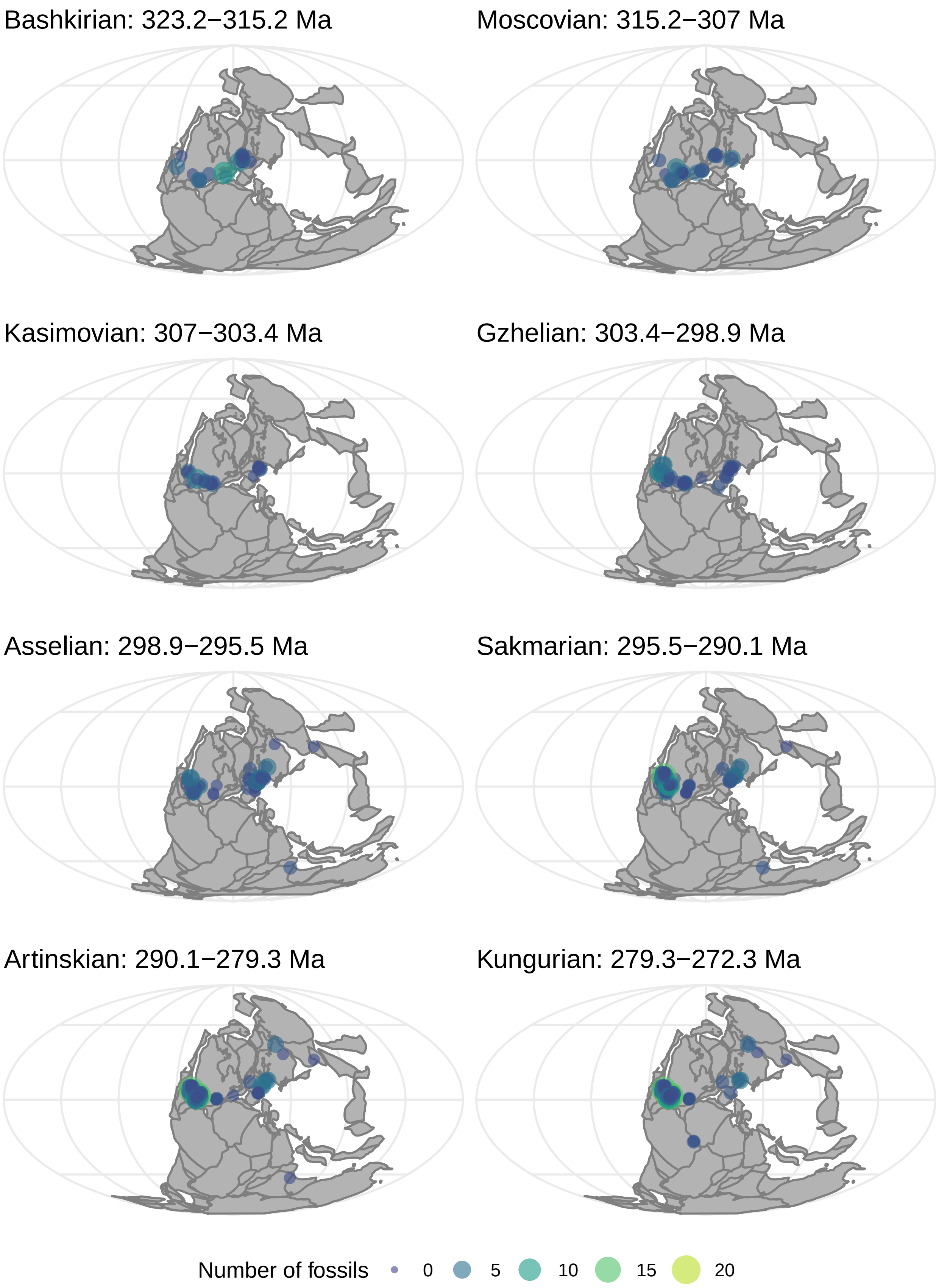

### Supplementary Figure 2

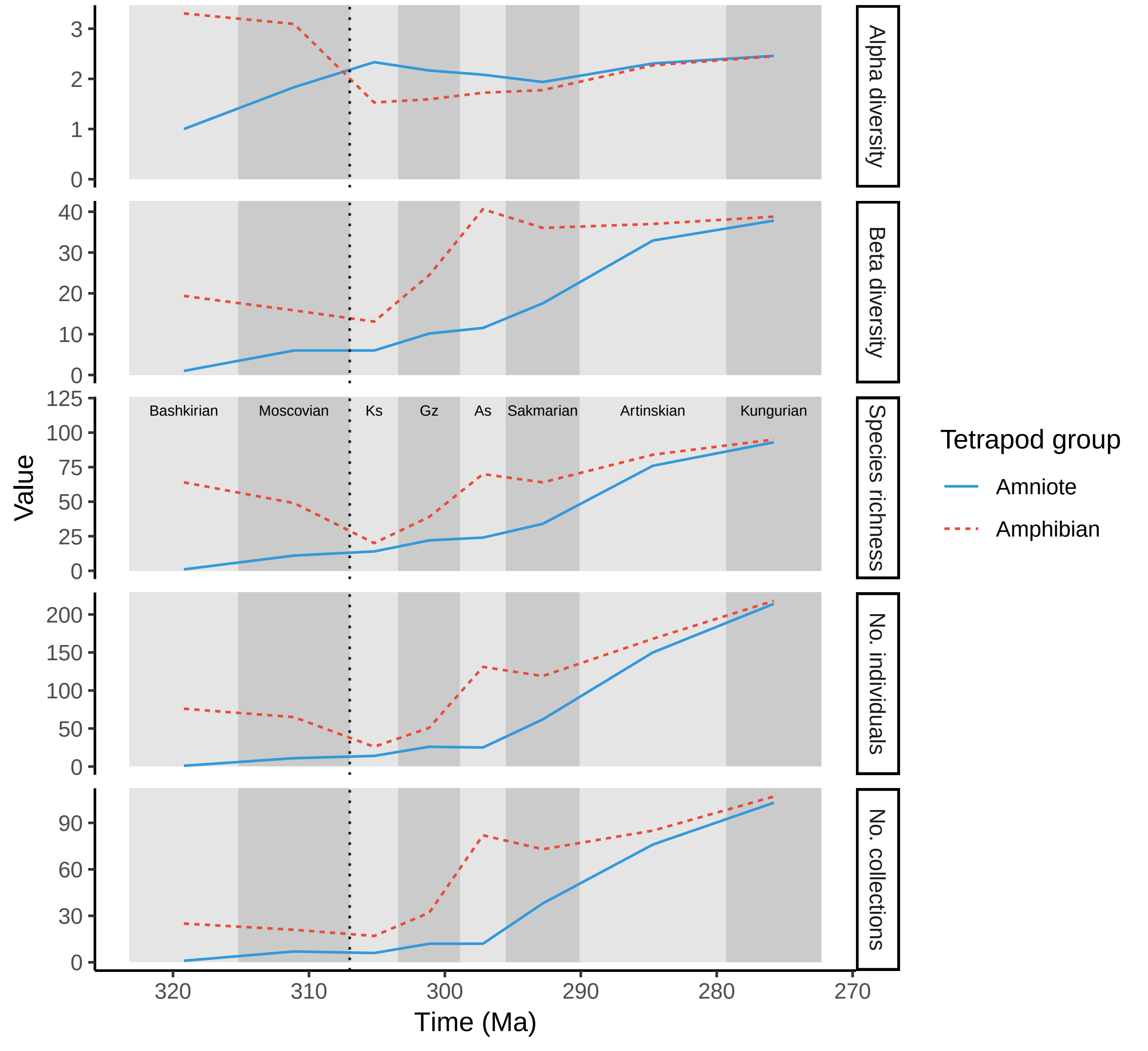

### Supplementary Figure 3

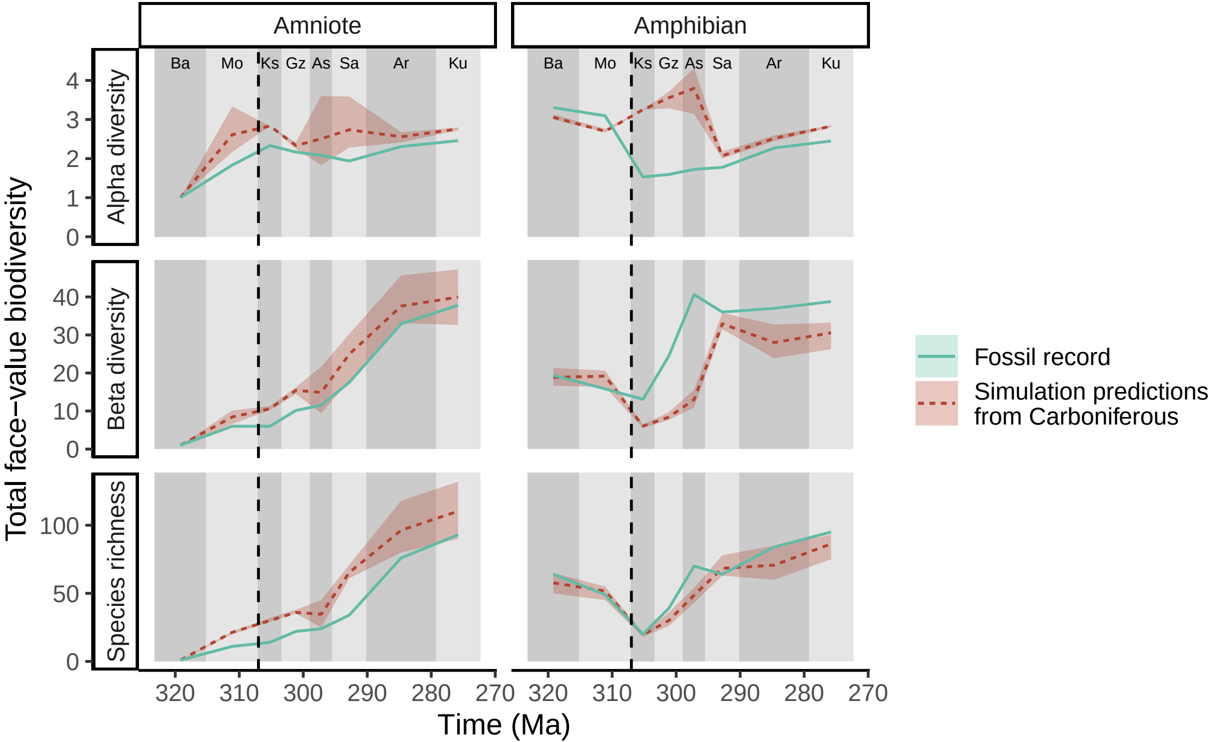

### Supplementary Figure 4

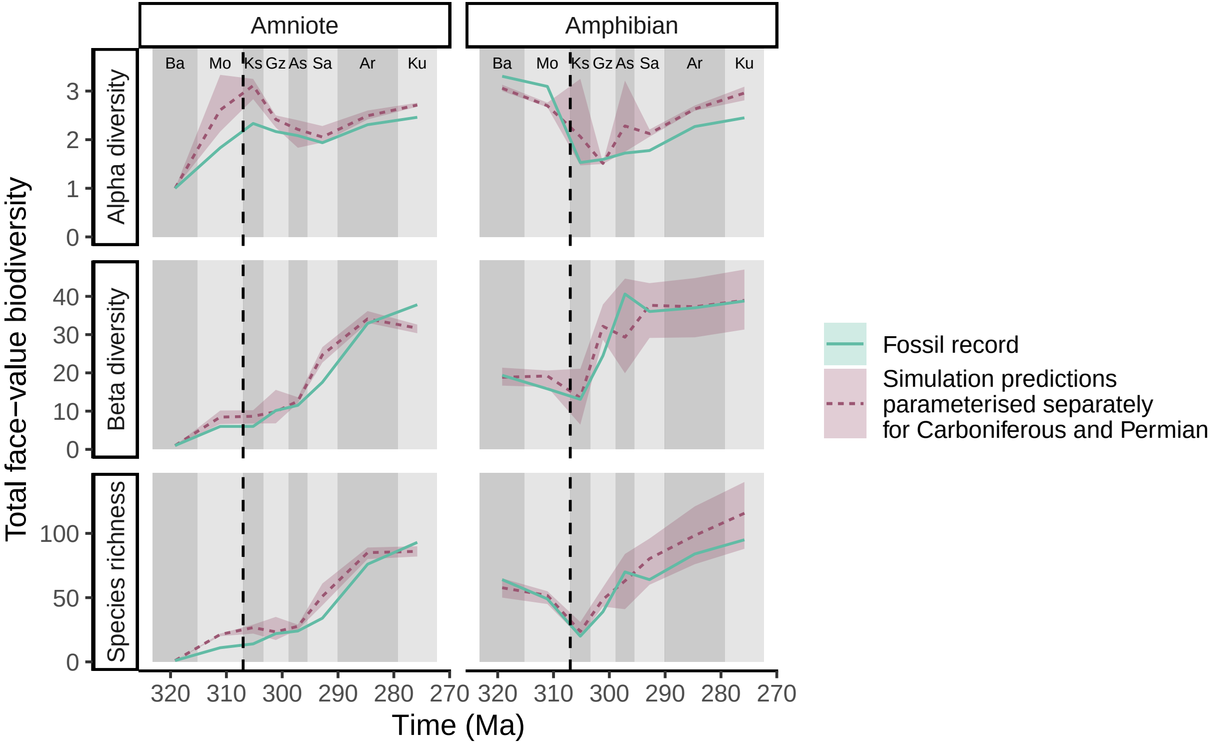

### Supplementary Figure 5

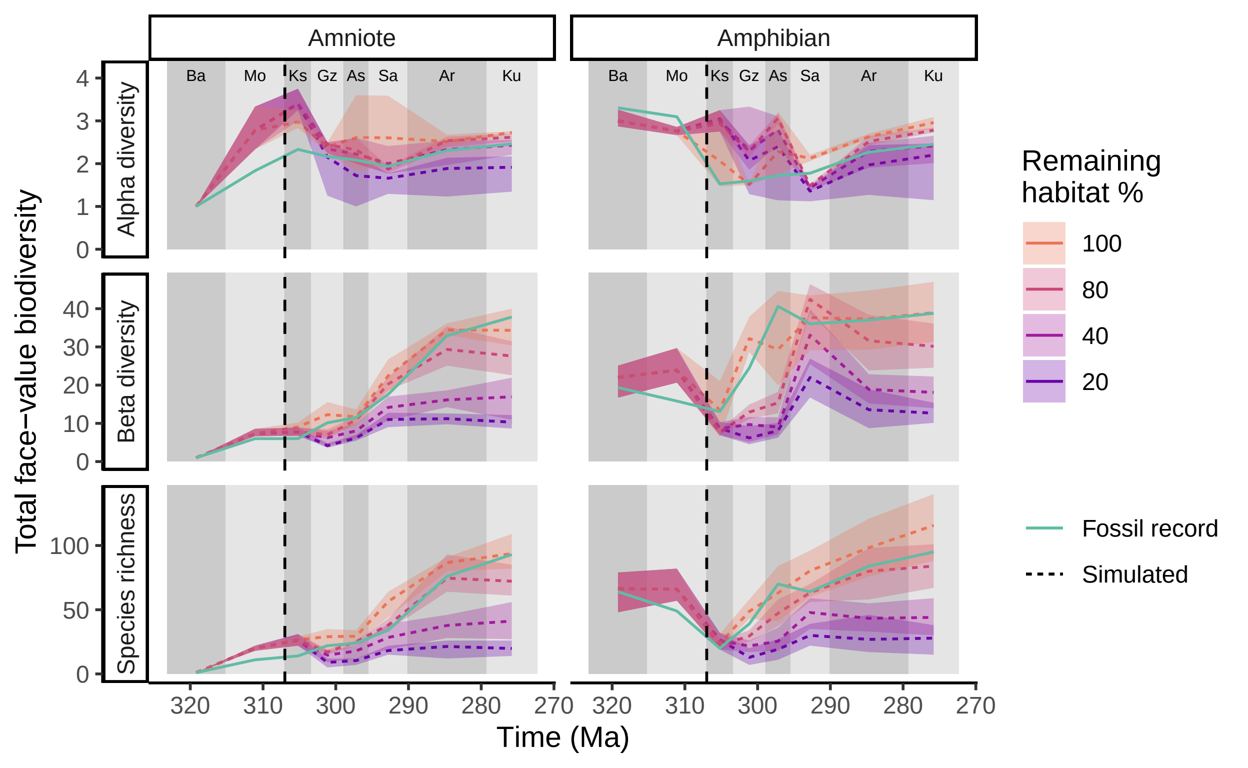
